# Metacontrast masking as a measure of change detection: Children with poor reading fluency show impaired change detection for both letters and shapes

**DOI:** 10.1101/697870

**Authors:** David Crewther, Jacqueline Rutkowski, Sheila Crewther

**Affiliations:** Brain Sciences Institute, Swinburne University of Technology, Victoria 3122, Australia; School of Psychological Science, La Trobe University, Victoria 3086, Australia

## Abstract

Developmental dyslexia, a specific learning difficulty in reading, manifests as effortful decoding of words and as such is commonly associated with reduced phonemic awareness. However, its underlying cause remains elusive, with magnocellular visual processing, temporal auditory processing, visual attentional deficits and cerebellar dysfunction all gaining some traction. More recent theories have concerned visual attention span, measuring the parallel attentive capacity of the sensory visual system. However the VA span task as implemented requires reports, both conscious recall and recognition of letters, that activate many cortical areas beyond sensory visual cortex. Change detection, in contrast, does not require the conscious recognition of items, but simply awareness that the stimulus has changed, or not, again testing visual attention in a parallel fashion, but avoiding the complications of higher order cognitive processes. Thus, we investigated change detection in 33 good and poor readers with ages of around 10 yr, using a gap paradigm. Groups of 4 letters or 4 shapes were presented for a fixed time (0.7 s), followed after a 0.25 s gap, by a second similar group, each item surrounded by an annular frame filled with dynamic random noise of variable contrast. Detection performance was manipulated by varying the contrast of these meta-contrast mask frames, yielding a threshold contrast of the frames at which participants could just detect change. In two separate experiments, letters and rectangular shapes were used as target items, in order to test whether previous findings of superior change detection in good compared with poor readers was a result of greater automaticity in letter recognition of the good readers. The results indicate that the good readers were able to detect change at higher levels of masking distraction for both the letter and shape targets, indicating that this difference is not specifically related to to the training of graphemic or lexical information but more likely reflects a difference in alerting or pre-recognition stages of visual processing. Together, the results provide further support of the notion that there is a low level attentional performance difference between dyslexic and normal reading children. Thus, the results further bring transient spatial attention directly into the spotlight as an ability critical for learning to read.

## Introduction

Developmental Dyslexia (DD) is an impairment in the acquisition of literacy skills despite normal intelligence, socioeconomic and educational circumstances, as defined under DSM-IV-TR (American Psychiatric Association, 2000), but positioned with specific learning disorder in DSM-5 (American Psychiatric Association, 2013). It is estimated to affect approximately 5-10% of school-aged children (Habib, 2000). These deficits tend to persist through to adulthood. Children with DD are reported to show deficits in many areas of perception, motor skills and learning but the primary deficit has long been considered to involve phonological processing (reviewed Bishop and Snowling (2004), Snowling and Hulme (2012)). Deficits in rapid temporal auditory processing have also been hypothesised as the basis of dyslexia by Merzenich et al. (1996), as dyslexics have difficulty in rapidly processing sequential information. Indeed, rapid temporal dot counting has been shown to be more difficult for children with dyslexia than spatial dot counting (Eden et al., 1995). Other multisensory processing deficits have been demonstrated with both auditory and visual gap detection impaired in children with dyslexia compared with normal reading children of the same age (Van Ingelghem et al., 2001) indicate the possibility of a general temporal processing deficit in children with dyslexia. More recent experiments employing stimuli designed to probe magnocellular activity have also identified perceptual speed as a common factor accounting for variance in reading ability (McLean et al., 2011).

Competing visual hypotheses also exist for the underlying mechanism of dyslexia. For example, dysfunction in the magnocellular visual pathway (Stein and Walsh, 1997;Habib, 2000;Gori et al., 2016) has been proposed on the basis of reduced motion and motion coherence sensitivity (Cornelissen et al., 1995;Talcott et al., 2000), and reduced brain activation in Area V5/MT+ (Eden et al., 1996) to moving stimuli with Demb et al. (1998) showing a 3-way interaction between brain activiation in MT+, thresholds for speed discrimination and reading speed.

Interest has also grown in abnormalities in transient attention as a basis for autistic perception. Indeed deficiencies in visual attention as the basis for dyslexia are supported by reports of slower serial search speeds and lowered search performance (Eskenazi and Diamond, 1983;Casco and Prunetti, 1996). Early work investigating the effect of backward masking (Grosser and Trzeciak, 1981) on the recognition of briefly presented single letters, demonstrated poorer performance in a group of dyslexic children compared with normals. Facoetti and Molteni (2001) showed that dyslexia is associated with an alteration in the gradient of attention with consequent abnormalities in engagement and disengagament (Facoetti et al., 2008). Visual attention span (VA) was introduced as a means of measuring the proportion of distinct visual elements that can be correctly processed in parallel in a briefly presented multi-element array (Bosse et al., 2007). Bosse et al. proposed that a deficit in VA span might contribute to developmental dyslexia, independently of abnormalities in phonological processing. This was supported through studies indicating a visual rather than verbal modality (Lobier et al., 2012). Also, fMRI studies using categorization of alphanumeric and non-alphanumeric character strings showed differential fMRI activation in the superior parietal lobule between participants with and without dyslexia (Lobier et al., 2014).

Other aspects of attentional processing, such as ability to detect change in the environment rapidly (change detection), have received relatively less attention, despite strong intergroup differences. Rutkowski et al. (2003) using a gap-paradigm change detection task, involving a stimulus array of 4 letters in circular place holders, showed that a much longer presentation time (an inspection time) for the initial group of letters was needed in order to just detect change, comparing dyslexic children with normal readers. There are some similarities between change detection and visual attention span, centred around the fact that for both tasks, several objects have to be processed in parallel. A key difference is that with change detection, the target objects do not have to be brought fully to conscious awareness in order to report change at better than chance level. VA span, on the other hand, requires identification and hence conscious perception of the objects. While change detection relies on the integration of the initial items in terms of depth of recognition, it is possible that the ability to detect change depends on low level attentional functions related to the sudden reappearance of the second group of items following the gap. If misdirection of attention provided by the place-holders around each object provides a masking signal, then this can potentially be used to measure change detection ability, independent of integration times for recognition of the initial display in the gap paradigm stmuli but rather as a function of ability to filter salient information by inhibiting distracting visual stimuli.

Thus we proposed to further investigate masking as a distracter in change detection, providing a novel thresholding mechanism using dynamic random noise of variable contrast in annular place holders as a variable meta-contrast masking of the second presentation stimulus of the gap change detection paradigm. This method of measuring change detection sensitivity (Rutkowski et al., 2003) was then used across letter and shape targets to establish whether previously reported change detection differences could be more attributable to better training in letter recognition in good readers rather than impairment in visual processing in those with dyslexia. The shape stimuli were used to test whether good readers, when presented with novel, untrained stimuli under similar task conditions, would perform any better than poor readers.

## Methods

### Participants

Thirty-three children were recruited from two non-selective state primary schools and from a summer camp for children with learning difficulties. Reading ability was assessed by trained experimenters using the The Neale Analysis of Reading Test (Neale, 1999). A reading age was established for each participant on the basis of the published norms. Assignment of participants to good and poor reading groups was on the basis of chronological age minus reading age

Two similar change detection tasks were carried out – one using detection of changes in letter identity (Expt 1a) and the other using change in orientation or colour of rectangular shapes (Expt 1b). Delay in the implementation of testing for Expt 1b resulted in overlapping but not identical populations for the two experiments.

**Table 1.** Participant Age and Reading Profiles for Experiments 1a and 1b.

| Stimulus | Reading Group | Chronological Age | Age Range | Reading Age | N |
| --- | --- | --- | --- | --- | --- |
| Letter | Poor | 10.9±0.6 | 7.0-14.2 | 7.7± 0.5 | 14 |
|  | Good | 9.2±0.3 | 5.7-12.2 | 11.3±0.5 | 17 |
| Shape | Poor | 9.0±0.9 | 7.0-13.6 | 7.4±0.7 | 7 |
|  | Good | 9.2±0.4 | 5.7-12.2 | 11.4±0.5 | 16 |

### Change Detection for Letter Stimuli

In this experiment, a fixation cross appeared for a period of 1 sec, prior to the appearance of the first stimulus, which consisted of four upper case letters, each subtending a visual angle of 1° high x 0.8° wide and presented at an eccentricity of 2° for a participant seated 57 cm from the computer monitor (see Figure 1a). The presentation time for the first stimulus was fixed at 0.7 sec, followed by a fixed gap of 0.25 sec (with only the fixation cross showing). The second display was as for the first display (with a probablity of 50% that one item was changed), except that each letter was surrounded by an annulus of width 0.2 subtending 2° at the eye filled with dynamic random noise (2 pixel), the contrast of which could be varied from 0 to 1. The second stimulus remained on screen until a button was pressed. Stimuli were created and presented using VPixx (www.vpixx.com) on an iMac computer (see Fig 1a). Participants were required (in a two alternative forced choice) to press the keyboard right arrow if they thought change had occurred and keyboard left arrow if they thought the two displays were identical. Convergence to threshold for the dynamic random dot noise was under the control of a PEST model. Participants performed 40 trials.

**Figure 1:**
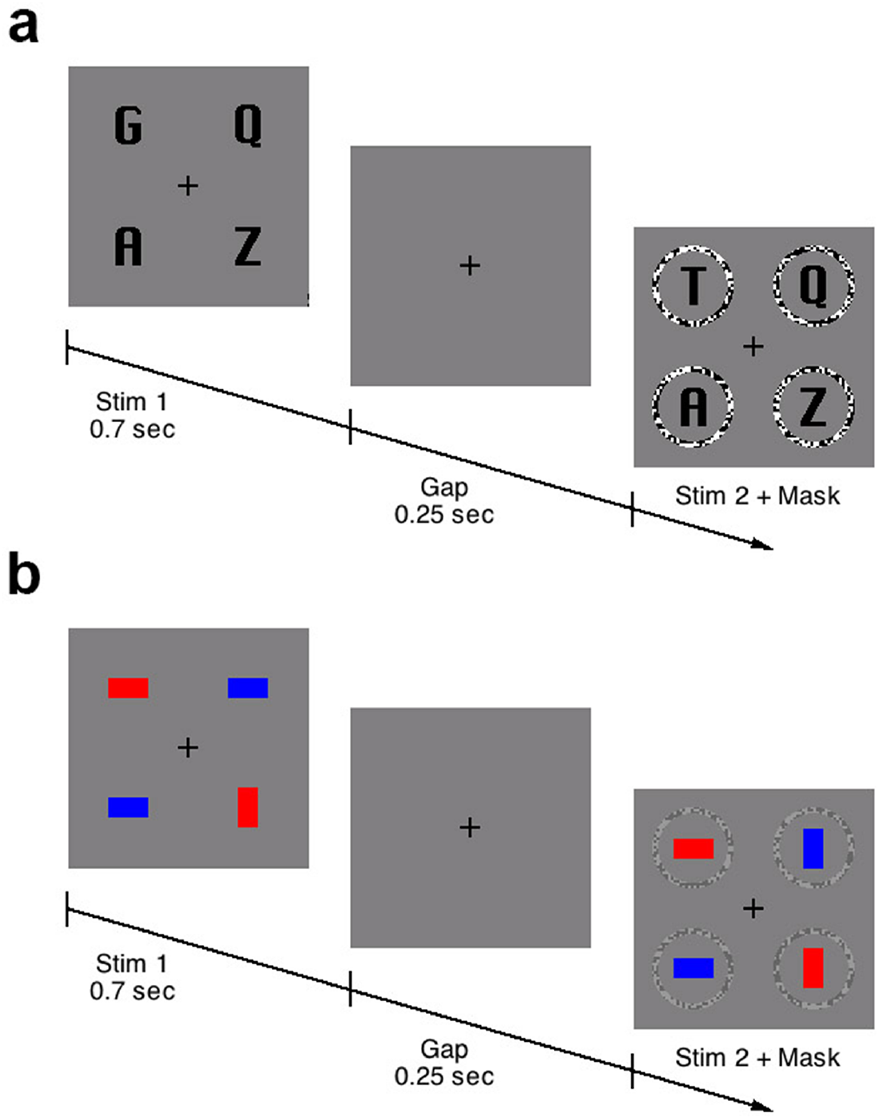
Experimental stimuli used in experimentas 1a and 1b. Both employed annular place holders around the 4 stimuli of the second presentation, with variable contrast random dynamic noise (shown at high contrast for the letter stimuli in Fig 1a, and low contrast in the shape stimuli for Fig 1b).

### Change Detection for Shape Stimuli

The shape experiment was identical to the letter experiment, except that shapes were substituted for letters. The shapes were rectangles oriented either horizontally or vertically and coloured either red or blue (see Fig. 1b). With 50% probability one of the shapes could change its orientation, its colour, or both, with conditions randomised. The shapes subtended 1° x 0.5° while the circular place holders subtended 2° at the eye of the participant seated 57 cm from the computer monitor. Stimuli were created and presented using VPixx on an iMac computer. Background luminance for both experiments was 50 cd/m^2^.

## Results

### Letters

Good readers performed signficantly better than poor readers on the change detection for letters task. This can be seen in Figure 2 which shows the mean threshold contrast (with standard errors) for the masking annuli. As the contrast of the flickering mask increases it becomes harder to detect change, and hence the contrast threshold represents a measure of change detection ability. Analysis of variance (ANOVA) was carried out on the threshold masking contrast compared between reading groups, finding a significant main effect of Reading Group (*F*(1, 29) = 8.35, *p* = 0.007).

**Figure 2:**
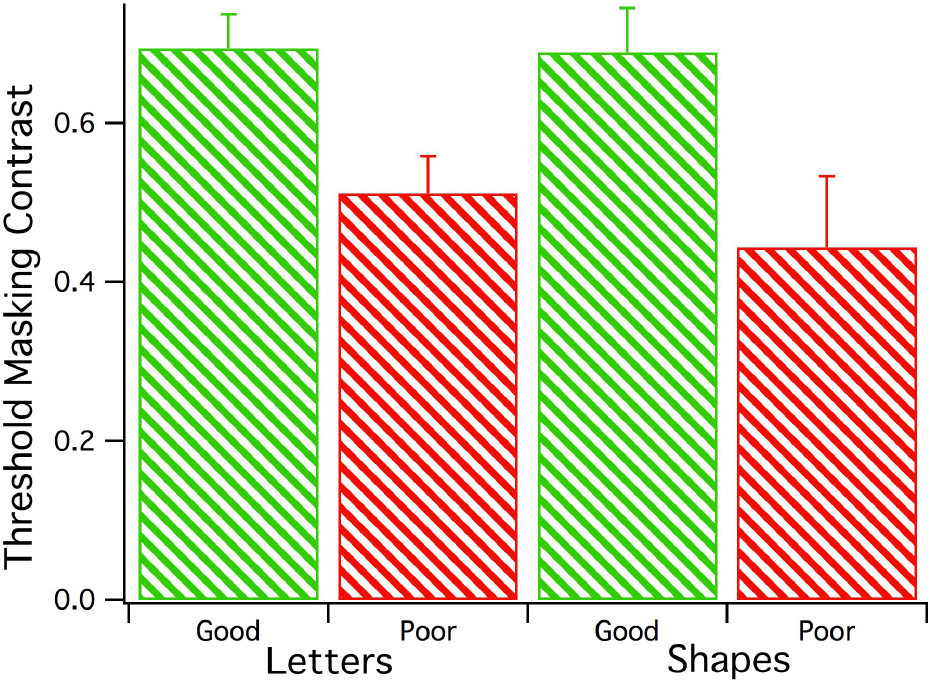
Threshold masking contrasts for letter and shape stimuli, for which groups of good and poor readers (GR, PR) could just detect change. The data are presented as means (with standard errors). Contrast thresholds for both letter and shape stimuli were higher for Good *cf* Poor reader groups.

### Experiment 1b: Shapes

Good readers performed better than Poor readers on the change detection for shapes as well (see Figure 2). Again, higher contrast indicates a better ability to withstand the distraction caused by the appearance of the rings. Despite the smaller population size of Poor readers in the Shapes experiment, the effect of reading group was again significant *F*(1, 22) = 5.75, *p* = 0.02).

The similarity in threshold results for the two tasks suggests the degree of correlation be investigated. Taken across all readers, the correlation between threholds for letters and shapes was significant (*F*(1,20) = 4.77, *p*=.04, *Rsq*=.19).

## Discussion

The results of these psychophysical experiments validate the use of variable masking as a measure of change detection, as the results replicate those established using different methods of measurement (titrating the initial display presentation time to estimate detection threshold) (Rutkowski et al., 2003). The results of this experiment reinforce the idea (Rutkowski et al., 2003) that the sudden appearance of the second stimulus is likely to be the factor that controls change blindness, for this type (display – gap – display) of change experiment.

The metacontrast masking method has the advantage that it opens up other possibilities for investigating the role of attention in change detection, particularly as it applies to the mechanisms of dyslexia. It has the benefit that a wide dynamic range of distraction is available – especially using the random dot noise which is qualitatively much more effective at high contrast than a static noise ring of the same geometrical dimensions and contrast. The fact that mask contrast can be used in such a controllable fashion to modify detection rate also suggests that the sudden appearance of the second stimulus (see Fig.1) may interfere with change detection precisely because attention is distributed across the field of possible targets. Indeed it would fit with better change detection for targets in upper visual field compared with targets in lower visual field (Rutkowski et al., 2003), in the following way. Primates including human appear to have a relative magnocellular concentration in lower compared with uper visual field (Previc, 1990;Pitzalis et al., 2012) in extrastriate cortical areas. This lower field bias for M cell activity has also been recently shown in multifocal VEP in children (Crewther et al., 2019). This bias, together with the contribution of M cells to transient attention make it likely that distraction from the flickering meta-contrast masks is likely to be stronger in lower visual field. This would cohere with the upper visual field change detection advantage seen with change detection stimuli surrounded by masks (Rutkowski et al., 2003).

A criticism of the interpretation of studies in dyslexia that use letters as stimuli is that letter processing by poor readers may be deficient because they have generally read less and practised letter recognition less. The discussion on both sides is well represented in Nature Reviews Neuroscience (Goswami, 2015b;a;Lobier and Valdois, 2015). This potential criticism provides the rationale for examining change detection using shapes. The second experiment, despite being less statistically powered, due to a smaller PR population, showed that when geometrical shapes were used as targets, change detection performance was still inferior for poor readers compared with good readers.

The significance of the difference in change detection between good fluent and poor nonfluent readers suggests that rapidity of object processing might play an important role in the development of fluent reading (see (Grosser and Trzeciak, 1981), irrespective of need for phonological awareness of various letters and words. This is borne out by recent experiments associating perceptual speed with reading ability (Hecht et al., 2004;Thomson et al., 2006;McLean et al., 2011). The current findings are hard to reconcile with Goswami (2015a) because while letters may have been learned better by those with good reading skills, the spatial arrangement of the change detection task is something that has not been trained, and yet the change detection performance on the shapes task was similarly impaired in those with poor reading skills. More likely as an an explanation is that the good readers exhibit better attentional control over low-level noise and distraction, irrespective of the targets.

## Acknowledgements

The authors would like to acknowledge support from the philanthropic Andrew Fildes Foundation for sponsoring summer camps, the Department of Education and Training Victoria, and an Australian Research Council Discovery Project.

## Notes

#### Summary of Updates

Figure 2 label added. Discussion clarified. Abstract revised for clarity. Typographical corrections made.

